# “Reassessing the origins of pathogenicity in *Candida auris* and relatives through phylogenomic analysis”

**DOI:** 10.1101/2023.04.13.536682

**Authors:** Kyle S. Schutz, Tina Melie, Stacey D. Smith, C. Alisha Quandt

## Abstract

Emerging fungal pathogens commonly originate from benign or non-pathogenic strains living in the natural environment. Assessing the evolutionary relationships between pathogenic and non-pathogenic species is one approach for tracing the origins of pathogenicity across species. The recently emerged human pathogen, *Candida auris* belongs to the *Candida/Clavispora* clade, a diverse group of 45 yeast species including human pathogens and environmental saprobes. *C. auris* is believed to have originated in the environment and recently transitioned to a human pathogen. We present a phylogenomic analysis of this clade aimed at testing for patterns implicated in the emergence of pathogenicity using an expanded sample of non-pathogenic strains and species. To build a robust framework for investigating these relationships, we developed a whole-genome sequence dataset of 108 isolates representing 18 species, including 4 newly sequenced species and 18 environmentally isolated strains. Our phylogeny, based on 619 orthologous genes, shows environmentally isolated species and strains interspersed with clinically isolated counterparts, rejecting the hypothesis of a single origin of pathogenicity within the lineage containing *C. auris* and its closest relatives. Our findings highlight the breadth of environments these yeasts inhabit, and imply, concerningly, that known pathogens could just as easily live outside the human body in diverse natural environments. Based on this result, we suggest that surveillance aimed at detecting emerging pathogens should expand to related environmentally-derived fungi with pathogenic potential.

**AUTHOR SUMMARY:** The rapid rise in the number of fungal pathogens over the past few decades has been linked to climate change, globalization, intensive farming practices, and an increase in immunocompromised individuals. *Candida auris* is an example of a recently emerged fungal pathogen capable of causing severe disease and large outbreaks in vulnerable patient populations. The evolutionary origins of *C. auris* are poorly understood, however, they are essential to understanding how and when this pathogen emerged. In this study, we investigated relationships between a sample of pathogenic and non-pathogenic strains and species in the *Candida/Clavispora* clade, a group of 45 yeast species including human pathogens (including *C. auris*) and environmental saprobes. We used these relationships to test for patterns that might support differing pathogen emergence hypotheses. We found that the relationships between pathogens and non-pathogens suggest many transitions between humans and other environments, rather than a single origin of pathogenicity. It seems plausible that these pathogens, often found in harsh environmental conditions such as seawater, already possessed traits that make them suitable human pathogens, which are perpetuated by increased at-risk patient populations. We should, therefore, be vigilant in our surveillance for clinical isolation of yeasts belonging to this clade from humans.

## INTRODUCTION

The emergence of new fungal pathogens poses a significant threat to human and animal health, as well as ecosystem health. Human fungal pathogens cause a range of diseases, from minor skin infections to life-threatening systemic mycoses. Systemic mycoses often have limited treatment options available and have the highest impact on individuals without access to health services, often becoming progressively worse if left untreated [1]. Humans are increasingly affected by fungal pathogens and opportunists, with an estimated 1.5 million deaths related to fungal infections reported each year [2,3]. The rapid rise of fungal pathogens over the last 30 years has been linked to climate change, globalization, intensive farming practices, and the increase in the number of immunocompromised individuals [4,5,6]. Emerging fungal pathogens can originate in different ways but commonly have evolutionary histories in benign or non-pathogenic species living in the natural environment [7]. It is often unclear how human infection enhances their fitness and becomes part of their natural lifestyle [8]. Delineating the evolutionary relationships between pathogenic and non-pathogenic species is a clear first step in assessing how virulence emerges and evolves across species [9].

While earlier research relied on marker genes for estimating relationships among fungal pathogens and their relatives [10], studies are increasingly moving to the use of whole genome sequence data [11], resulting in greater clarity and confidence in phylogenetic estimates. These phylogenetic advances have often revealed that traditional taxonomic designations, relied upon for classifying and comparing fungi in clinical contexts, may not correspond to natural (monophyletic) groups [10]. A prime example is the highly polyphyletic budding yeast genus *Candida*, which is represented by at least 30 distinct lineages scattered across the subphylum Saccharomycotina [10]. This genus includes 31 species known to cause disease in humans, including *Candida albicans, Candida glabrata, Candida tropicalis, Candida parapsilosis*, and *Candida krusei*, which together account for roughly 90% of *Candida* infections [12,13]. However, the most concerning and enigmatic yeast in this genus may be *C. auris*, a deadly, multi-drug-resistant pathogen that has caused large outbreaks in hospital settings and high mortality in vulnerable populations [14,15]. *Candida auris* belongs to the *Candida/Clavispora* clade (*Metschnikowiaceae*), a group of 45 yeast species including human pathogens and environmental saprobes [10,11,16,17,18,19,20]. Nevertheless, the exact placement of *C. auris* within the clade (and thus the identity of its closest relatives) remains uncertain as previous phylogenetic studies have recovered conflicting results. Moreover, while whole genome data has been available for *C. auris* for several years [21], we lack such genomic data for related non-pathogenic species, precluding more robust phylogenetic inference within the *Candida/Clavispora* clade [22].

The wide range of environments from which members of the *Candida/Clavispora* clade have been recovered suggests broad ecological niches, even for the pathogenic species [23]. The non-pathogenic species have been isolated from diverse sources and locales, including flower nectar from flowers of *Ruellia* spp. in India [24] and a cockroach’s gut from Barro Colorado Island, Panama [25]. Only six of the 45 described species have been isolated from human sources consistently (*C. auris, C. haemulonii, C. pseudohaemulonii, C. duobushaemulonii, C. vulturna*, and *Clavispora lusitaniae*) [21] and all of these (apart from *C. pseudohaemulonii*) have been recovered from environments outside of clinical settings. For example, *C. haemulonii* was first isolated in Florida in 1962 from the gut of a blue-striped grunt (*Haemulon sciurus*), and it has since been isolated from seawater in Portugal and other marine sources [16,26,27]. Similarly, *C. duobushaemulonii* was first isolated from insect frass in Germany [16] and *C. vulturna* from flowers in the Philippines [20,28]. *C. auris*, by contrast, was first identified in clinical samples and only recently isolated from salt marshes and beaches in the Andaman and Nicobar Islands in Southeast Asia [29]. The diverse isolation sources of these pathogens, from seawater to blood, raise questions about their metabolic versatility and whether it played a role in their success in overcoming the barriers to surviving the human body.

Broadly, fungi must overcome many physiological challenges to living in the human body, including growth at elevated temperatures and high saline concentrations, while simultaneously evading the human immune response [30]. Specific virulence factors, such as anti-fungal resistance profiles, physiological traits, and epidemiology vary among pathogens in the *Candida/Clavispora* clade. *C. auris* is a unique pathogen known for its thermotolerance (growth above 42°C), multi-drug resistant phenotype, persistence on abiotic surfaces, and ability to transmit from patient to patient via asymptomatic skin carriage [27]. These traits vary across *C. auris* strains, which form four genetically distinct sub-clades and vary in their propensity for invasive and superficial infections [31]. Pathogens in the closely related *C. haemulonii* species complex (which typically includes *C. duobushaemulonii, C. haemulonii, C. pseudohaemulonii*, and *C. vulturna*) are less thermotolerant but share resistance profiles with *C. auris*, such as resistance to amphotericin B and reduced susceptibility to azoles and echinocandins [16,21]. Due to growth restrictions at higher temperatures, these species tend to cause superficial infections, although invasive infections are increasingly reported [33]. Unlike *C. auris* and pathogens in the *C. haemulonii* complex, *Clavispora lusitaniae* seldom causes infection in humans and is typically described as an opportunistic pathogen, frequently in immunocompromised patients with comorbidities [33].

Given the sudden emergence of *C. auris* as a global pathogen and the diversity of isolation sources for it and other members of the *Candida/Clavispora* clade, several hypotheses have been put forward to explain the evolution of its pathogenicity. Many are rooted the endemic pathogen hypothesis, namely that a yeast exists in the environment but undergoes certain genetic changes that allow it to survive in the human body [7]. The Global Warming hypothesis specifically posits that *C. auris* transitioned from an environmental saprobe to a human pathogen due to adaptation to increasing ambient temperatures allowing for strains to overcome the thermal restriction zone [17, 29]. Other hypotheses suggest that these yeasts long existed as benign commensals of the human mycobiome but evolved pathogenicity due to changes in the host and host environment [27]. This hypothesis is embedded in the HIV/AIDS pandemic, where suddenly a large population of immunocompromised individuals was undergoing antifungal treatment [4]. An alternative to the endemic pathogen hypothesis is the novel pathogen hypothesis. This posits that yeasts which already possess traits that would make them suitable pathogens are being provided with a new host as humans encroach upon more territories and previously uninhabited environments [7].

In this study, we present a phylogenomic analysis of the *Candida/Clavispora* clade aimed at testing for patterns implicated by these differing emergence hypotheses by expanding the sampling of non-pathogenic strains and species. Specifically, we expect that if pathogens like *C. auris* began as environmental saprobes, the strains isolated from clinical environments will form a clade sister to or nested inside of non-pathogenic strains. To build a robust framework for investigating these relationships, we developed a whole-genome sequence dataset of 108 isolates representing 18 species, including 4 newly sequenced species and 18 environmentally isolated strains. Our findings provide a new evolutionary context for the emergence of the deadly human pathogen of global importance, *C. auris*, and other concerning human pathogens belonging to the *Candida/Clavispora* clade.

## RESULTS/DISCUSSION

### Diversifying Taxon Sampling in the *Candida/Clavispora* Clade

Species belonging to the *Candida/Clavispora* clade were identified through a review of existing phylogenetic estimates based on whole genome and marker gene sequence data, as well as historical taxonomic data from publicly available databases [11,18,19,21,27]. From this review, we estimated that 45 species comprise the *Candida/Clavispora* clade. Of those 45 species, however, 27 species did not have available raw whole genome sequence data or isolates. We identified 4 species with raw whole genome sequence data from GenBank (*Candida blattae, Candida intermedia, Candida oregonensis*, and *Candida thailandica*) and 5 species with isolates available for sequencing (*Candida dosseyi, Candida hainanesis, Candida heveicola, Candida mogii*, and *Candida ruelliae*) (Table 1). These 9 species were all isolated from environmental substrates and have not been described as human pathogens. The remaining 9 *Candida/Clavispora* species in our dataset have abundant isolates or sequence data available, although the majority of these have been isolated clinically. To diversify isolation source within our dataset, we identified environmentally isolated strains from known pathogenic species to include (Table 1). Although these isolation sources do not necessarily correspond to an ecological niche, they do represent conditions in which viable yeast cells were cultured. In total, our final dataset included 71 clinically isolated strains and 37 strains isolated from plants, non-human animals, or other environments (S1 Appendix, Fig 1). Based on broader phylogenies, we selected *Saccharomyces cerevisiae* and *Metschnikowia bicuspidata* as outgroups for our phylogeny. We rooted all trees on *S. cerevisiae*.

**Table 1:** Description of isolates whole genome sequenced in this study.

| Species | Strain | Collection Country | Substrate | Collect Date | CDS Region | Busco Score (%) | GC % | N50 | Assembly Size (Mb) |
| --- | --- | --- | --- | --- | --- | --- | --- | --- | --- |
| *C. dosseyi* [58] | CBS 10313 | USA | *Ululodes macleayanus* | 2005 | 6430 | 99.3 | 53 | 308944 | 15.5 |
| *C. duobushaemulonii* [16] | CBS 9754 | Germany | *Pyrrhocoris apterus* | 2012 | 5299 | 98.8 | 49.39 | 352276 | 15.2 |
| *C. haemulonii* [16] | CBS 5150 | Portugal | Seawater | 2012 | 5265 | 98.8 | 47.62 | 268833 | 15.2 |
| *C. hainanensis* [59] | CBS 10696 | China | Flower - *Magnoliaceae* | 2008 | 5195 | 99 | 52 | 393541 | 12.6 |
| *C. heveicola* [59] | CBS 7249 | China | Rubber Tree Sap | 2008 | 8246 | 99.2 | 49.22 | 550567 | 15.6 |
| *C. heveicola* [59] | CBS 10701 | China | Rubber Tree Sap | 2008 | 5292 | 99.1 | 47 | 268080 | 22.3 |
| *C. mogii* [60] | CBS 2032 | Japan | Fucho-Miso | 1967 | 4737 | 98.8 | 46 | 112524 | 12.9 |
| *C. ruelliae* [24] | CBS 10815 | India | Flower - *Ruelliacae* | 2012 | 5210 | 98.7 | 46 | 545101 | 14.2 |

**Fig 1:**
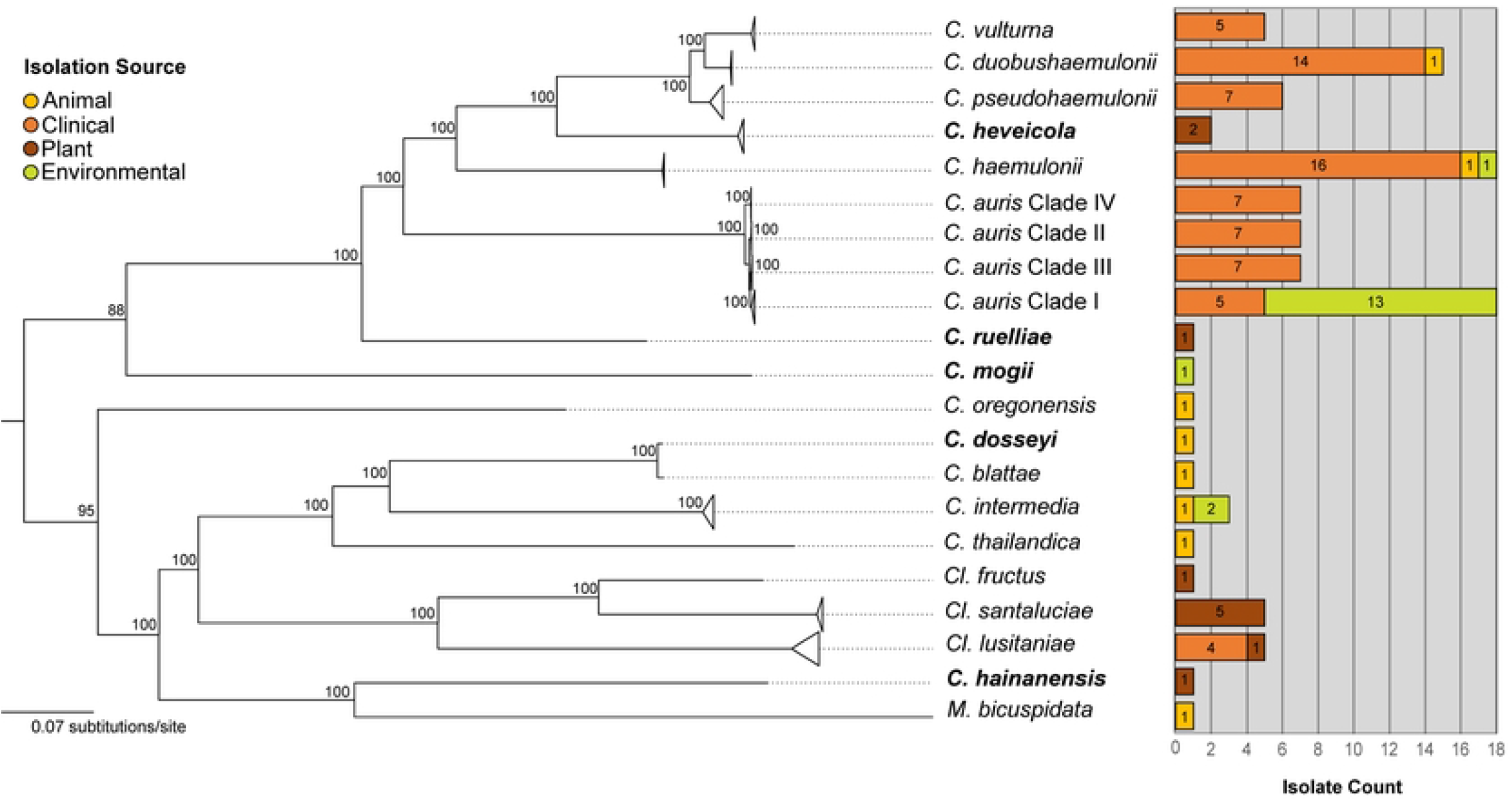
Multiple origins of pathogenic lifestyle in sampled *Candida/Clavispora* yeasts. A collapsed maximum likelihood tree rooted on *S. cerevisiae* (pruned) based on 619 orthologous genes from 108 isolates using a GTR+GAMMA model and 100 bootstrap replicates. Triangles represent a collapsed node. Bolded species were newly whole genome sequenced as part of this study. Bar graph represents the sampling diversity of isolation sources by category used in this dataset. An expanded version of this phylogeny can be found in Supplemental Fig 1.

### Phylogeny estimation for the *Candida/Clavispora* clade based on orthologous genes

From our 108 genome assemblies (82 *de novo* and 26 reference), we recovered 619 single-copy orthologous genes, similar to the number of shared core genes to other studies analyzing yeasts in this family [9]. Maximum likelihood (ML) phylogenetic inference based on the concatenated dataset showed high bootstrap support for relationships among species and clades (Fig 1). The branch lengths separating strains were orders of magnitude shorter than those separating species, suggesting recent divergence and few genetic differences among the strains. Given that such rapid speciation events can lead to discordant gene trees [34], we also estimated a species tree using multispecies coalescent methods that account for incomplete lineage sorting [35]. The species tree topology (Supplemental Fig 1) recovered the same relationships among the species as in the ML phylogeny but differed in some of the shallow relationships among the intraspecific strains (Supplemental Fig 1). Observing the same clades corresponding to species in both analyses strongly supports the current species delimitation in the *Candida/Clavispora* clade. Moreover, gene trees were generally concordant for these species-level clades (Supplemental Fig 2). While some intraspecific clades had strong agreement across genes (e.g., the clades within *C. auris*, Supplemental Fig 2), most shallow relationships showed conflict and uncertainty across gene trees (Supplemental Fig 2). This pattern is, again, consistent with the notion that the strains within species are only recently diverged, with short internodes leading to incomplete lineage sorting [36]. Many genes are also uninformative at the intraspecific level (gray portion of pies in Supplemental Fig 2), implying that they have little phylogenetically informative variation at that scale.

### Relationships between *Candida/Clavispora* species

We recovered many previously supported relationships but also discovered new relationships, particularly for the newly sampled taxa. The addition of *C. dosseyi, C. heveicola, C. hainanensis, C. mogii*, and *C. ruelliae* to the phylogenetic analyses helped resolve the uncertainty of their placement in the clade, as well as their relationships to known pathogenic relatives. *C. dosseyi* (isolated from an insect) is part of the *Clavispora* sub-clade, sister to *C. blattae* (also isolated from an insect) (Fig 1). These two species are nested within a larger clade with two other environmentally isolated species *Cl. intermedia* strains from sewage and wheat-straw hydrolysate, and a *C. thailandica* strain isolated from insect frass (Fig 1). These species are sister to *Cl. lusitaniae* (Fig 1). This topology is similar to a recent phylogenetic estimation of *Clavispora* species based on aligned proteins [18] but differs in the placement of *M. bicuspidata*, which here is sister to the newly added *C. hainanensis* (bootstrap support of 100% from the concatenated dataset; supported by 500/619 genes with 50% or higher bootstrap, Supplemental Fig 1 and 2).

*C. ruelliae*, isolated from flowers in India, is a yeast that demonstrates thermotolerance and the ability to form pseudohyphae, traits commonly associated with pathogenic potential [15]. *C. ruelliae* had been previously estimated to be part of the *C. haemulonii* complex species [15,20]. Prior to this study, a whole genome sequence had not been produced for the only known *C. ruelliae* isolate. In our topology, *C. ruelliae* is sister to both *C. auris* and the *C. haemulonii* complex species with 100% bootstrap in the concatenated analysis and 125 gene trees individually in support of this topology (Fig 1; Fig 2). *C. mogii*, isolated from Fucho-Miso soybean paste in 1962, appears to be sister to the *C. auris, C. haemulonii* complex species and *C. ruelliae* clade (Fig 1; Fig 2). Although this isolate was previously estimated to be distant from these yeasts based on marker gene sequencing data [20], the topology inferred here shows 88% bootstrap support and 82 gene trees in agreement with its placement.

**Fig 2:**
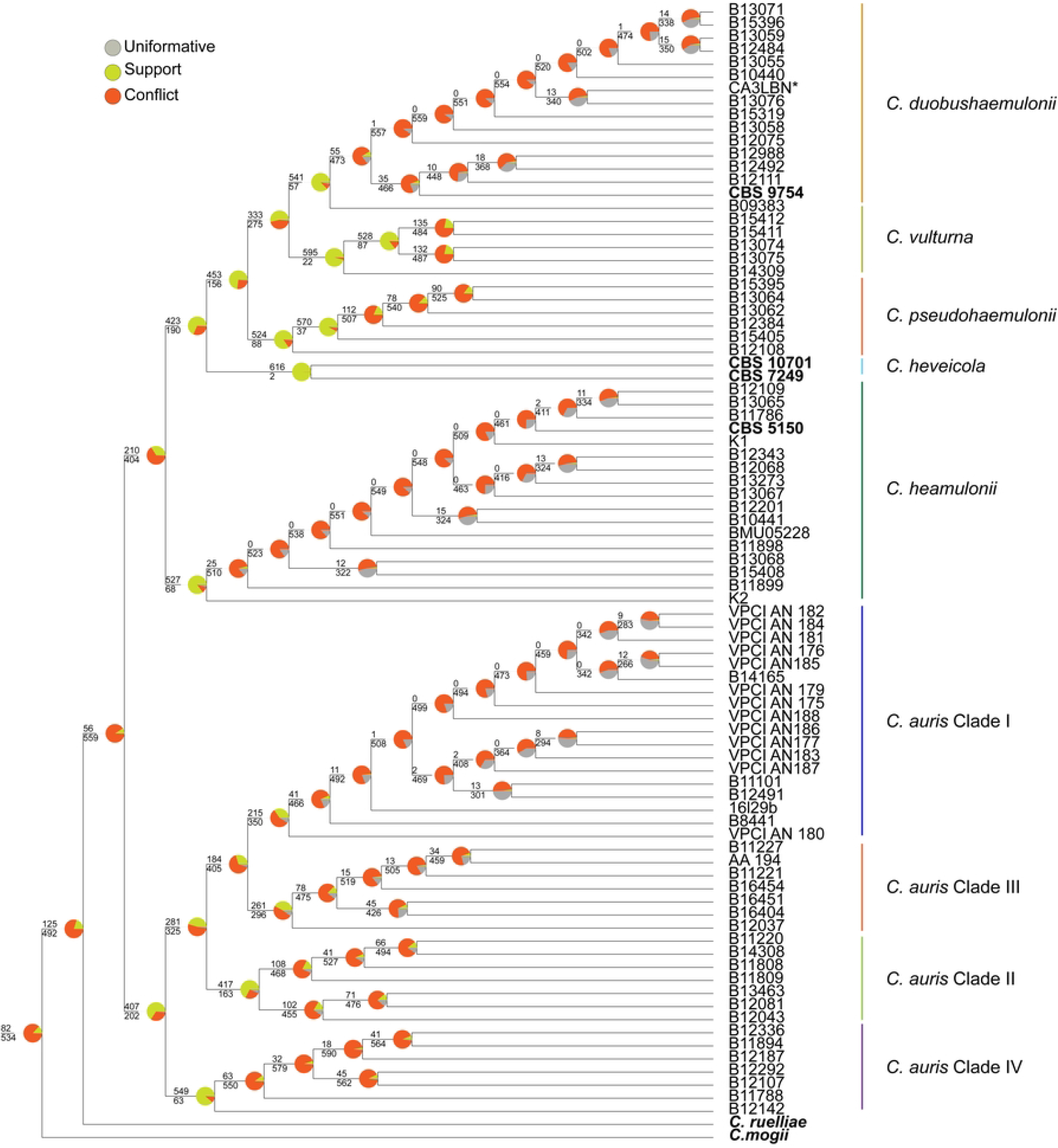
Recent divergences of *C. auris* and *C. haemulonii* show conflict among orthologous ML genes trees. A coalescent species tree estimation based on 619 ML gene trees in ASTRAL. Gene conflict was assessed by a conflict analysis using PhyParts. Green portion of the pie charts show the proportion of 619 ML gene trees that support on the presented topology with 50% or higher bootstrap. Red portion of the pie charts show ML gene trees that support an alternate topology with 50% or higher bootstrap, while the grey portion of the pie charts depict genes that are uninformative (< 50% bootstrap support).

### Assessing origins of pathogenicity *Candida/Clavispora* clade species

#### C. haemulonii *complex species are not exclusively pathogens*

The *C. haemulonii* complex species are often presented as a single clade of multi-drug resistant, human-pathogenic fungi sister to *C. auris*, an informal taxonomic designation often based on the inclusion of the following pathogenic species (*C. duobushaemulonii, C. haemulonii, C. pseudohaemulonii*, and *C. vulturna*) [20,37,38]. In this evolutionary scenario, these pathogenic species would all share a most recent common ancestor, pointing to a single origin of pathogenicity among these yeasts nested within the larger *Candida/Clavispora* clade. To test this hypothesis, we included genome-level data for environmentally isolated strains of species hypothesized to be part of this species complex. We included two strains of *C. heveicola*, isolated from rubber tree sap in China, as well as environmentally isolated strains of C. *duobushaemulonii* and *C. haemulonii* [20,24,26].

Our phylogenetic reappraisal of this species complex estimated that environmentally isolated *C. heveicola* is nested within the species complex and shares a most recent common ancestor with *C. haemulonii* and is sister to other *C. haemulonii* complex pathogenic yeasts (*C. vulturna, C. pseudohaemulonii and C. duobushaemulonii*) (Fig 1). There is high bootstrap support (100%) and 98% of the individual gene trees support this placement (Fig 1; Supplemental Fig 1). *C. heveicola* is the only species within this complex that has not been isolated from a human source, suggesting a single origin of pathogenicity is unlikely in the *C. haemulonii* complex and *C. auris* lineages. Additionally, we found that the environmentally isolated *C. duobushaemulonii and C. haemulonii* strains are interspersed within clinical strains (Fig 3a). A relatively old strain, *C. haemulonii* B10441 was isolated from the gut of a blue-striped grunt (*Haemulon sciurus*) in 1962 [26]. This strain clusters with a clinical strain isolated from blood in Panama in 2017. *C. haemulonii* CBS 5150 isolated from seawater in Portugal in 2017 is distantly related to strain B10441 but is sister to a clade of clinical isolates from Israel, Panama, and Venezuela (Fig 3a). Similarly, environmentally isolated *C. duobushaemulonii* is interspersed within the clinically isolated strains and is sister to strain B12988 isolated from a fingernail in North America (Supplemental Fig 2). Given the high gene tree conflict in this clade (Fig 2), we were interested to know if any of the 619 individual gene trees estimated a topology consistent with a single origin of pathogenicity within *C. haemulonii* and *C. duobushaemulonii*. We applied a topological filter in PAUP* v. 4.0a169 [39] to the 619 ML gene trees but did not recover any ML topologies that support environmentally isolated strains to form a single clade in both species.

**Fig 3:**
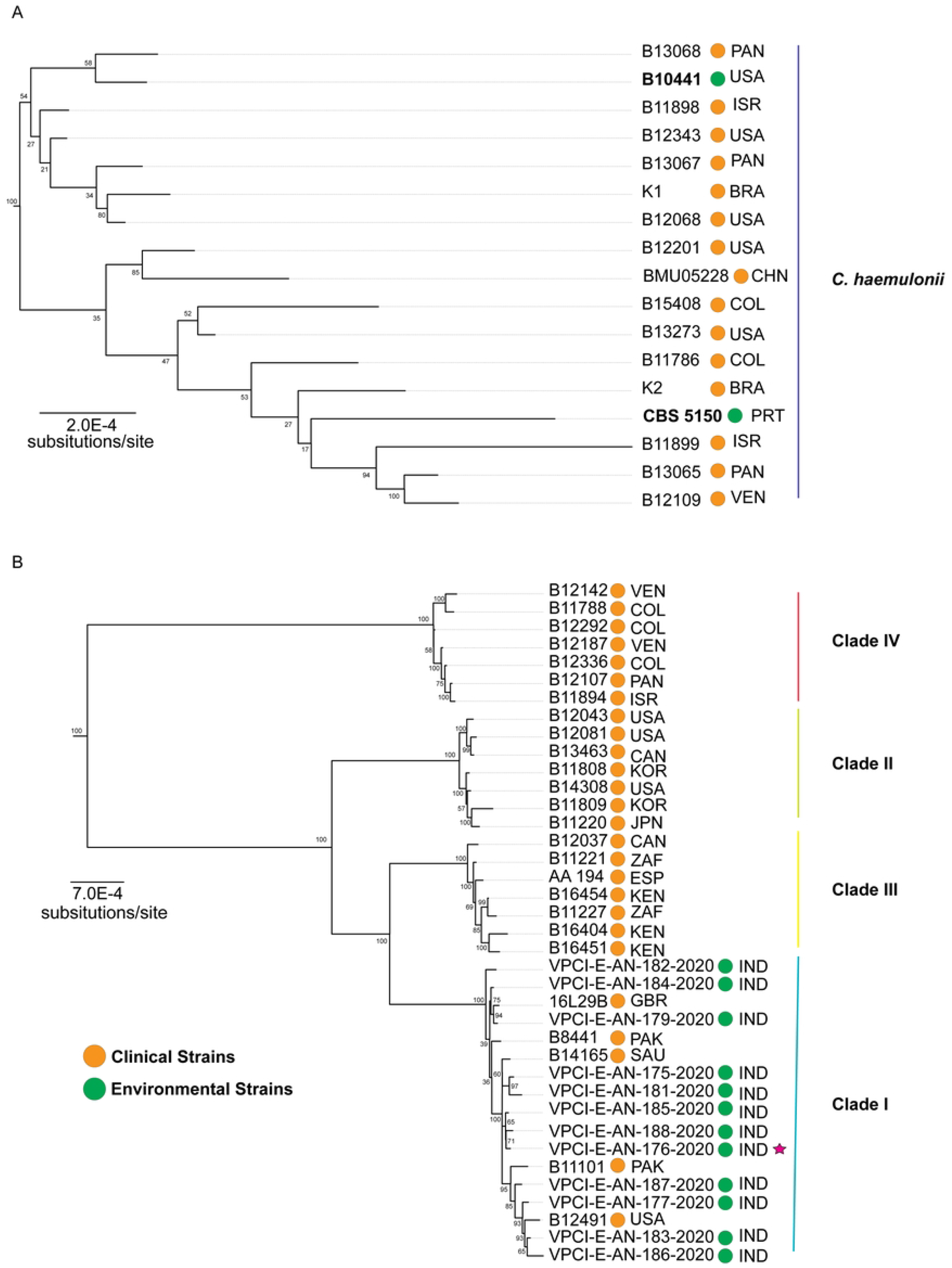
Strains isolated from non-clinical sources are not separate environmental lineages of *C. haemulonii* and *C. auris*. Maximum likelihood tree rooted on *S. cerevisiae* based on 619 orthologous genes using a GTR+GAMMA model and 100 bootstrap replicates. Three letters represent the country where the sample was collected. **A)** Zoomed in phylogeny showing sampled *C. haemulonii* strains. Green circles represent environmentally isolated strains. CBS 5150 was isolated from seawater in Portugal in 1972 and B10441 was isolated from blue-striped grunt in 1962. **B)** Zoomed in phylogeny showing sampled *C. auris* strains. Corresponding clade number was assigned based in NCBI designation. Environmentally isolated strains (green circles) represent isolates collected by Arora et al. 2021 in the South Andaman Islands. VPCI-E-AN-180 was pruned from the tree due to poor sequence quality. The red star denotes an antifungal susceptible, environmentally isolated *C. auris* strain (fluconazole MIC, 8 mg/liter; amphotericin B MIC, 1 mg/liter) [29].

In sum, *C. haemulonii* complex species do share a recent common ancestor with *C. auris*, but it is not exclusively a pathogenic species complex (Fig 1; Fig 3a). Additionally, at the strain level, there are no individual gene trees that support a topology in which environmentally isolated strains form a single clade. Taken together, it seems unlikely that these yeasts evolved from a most recent common ancestor that was strictly environmental and non-pathogenic. It is possible that addition of newly identified environmentally isolated strains of these species could clarify the topology by way of environmental sister groups or even a grade of environmental samples, which could support a transition from the natural environment to human, but even so, there would still be environmental strains nested within clinical isolates, consistent with escapes from human habitats or multiple environmental spillovers into human populations. Although it would be challenging to achieve sufficiently dense sampling to robustly estimate the number and direction of transitions, the intermixing of clinical and environmental isolates in our analyses highlights the breadth of environments these yeasts inhabit and suggests flexibility in their growth conditions.

#### *ML phylogeny captures the versatility of lifestyles in* C. auris

With our whole genome data, we estimated relationships between the newly identified environmental *C. auris* strains and clinical strains representing the four known sub-clades of *C. auris*. The isolation of *C. auris* from salt marshes and sandy beaches in the Nicobar and Andaman Islands marked the first time this species had been found in the natural environment, suggesting a possible environmental niche for the deadly pathogen which bolstered previous hypotheses about its ecological origins in hot, humid climates [17,29]. Similar to previous analyses, we recovered four, highly supported sub-clades of *C. auris*, which together form a clade sister to the *C. haemulonii* species complex [21, 31] (Fig 2a; S1Appendix). The branch subtending *C. auris*, is relatively long with many substitutions over time (Fig 1), while branches within the subclades have little agreement across gene trees (Fig 2) and are very short, consistent with previous intraspecific analyses of *C. auris* [21,31].

Based on these data, we found no genetic basis for a single origin of pathogenicity in *C. auris*, as the environmentally isolated strains from salt marshes were interspersed within the clinically isolated *C. auris* strains (Fig 3b). Environmentally isolated strains from the Andaman Islands do not form a genetically distinct clade from clinically isolated strains in Clade I. Given the high proportion of conflicting or uninformative gene trees at the terminal nodes in Clade I, we were also interested to know if any of the 619 individual gene trees presented a topology where environmentally isolated strains for a single clade (Fig 2). We applied a topological filter to the 619 ML gene trees but did not recover any gene trees that support environmentally isolated strains forming a single clade in Clade I.

Our topology captures the versatility of lifestyles in *C. auris*, from invasive pathogen to marine yeast. We did not find evidence that these environmentally isolated strains represent a separate lineage from the natural environment; instead, the isolation sources suggest that these yeasts may just as easily live in salt marshes as they could live on the human body. A recent study scanning metagenomic databases recovered marker-gene traces of *C. auris* and relatives in an even broader geographic range in samples from diverse environments, such as amphibian skin and soil [40]. Taken together, this evidence suggests that it will be challenging to isolate all of the ecological reservoirs of *C. auris* in natural environment, given that it can likely survive many harsh conditions. It is also unclear from these data, how these yeasts are growing and surviving outside the human body. Therefore, emphasis should be placed on studying their capability to survive harsh conditions and how that relates to their rapid success in the human body [8].

## CONCLUSION

The patterns of phylogenetic relationships recovered in our analyses suggest many transitions between humans and other environments. Using isolation source as a proxy for ecological breadth, we found that yeasts in the *Candida/Clavispora* clade likely have a high capacity for transition between many different environments. Specifically, we found that environmental isolates are commonly nested within clades of clinical isolates, as opposed to forming distinct clades sister to clinical isolates, underscoring that there is not a single divergence event that gave rise to a group comprised exclusively of human pathogens. This parallels the origins of pathogenicity in other clades of *Saccharomycotina*, where transitions of non-pathogens to the human pathogens have occurred independently at least five times [30]. In these *Candida/Clavispora* yeasts as in other *Saccharomycotina*, colonization of a human host could be a side effect of their physiological capability to survive in diverse, harsh conditions and not necessarily the gain of specialized virulence traits, which likely evolve due to selection pressure present during infection [8]. In addition to regulatory and coding sequence changes, these traits may be tied to changes in gene content, such as gene gain and loss as is the case for *C. albicans* [41], which we can now begin to explore with the growing body of genome assemblies.

This study provides a foundation for future hypothesis testing regarding the emergence of novel human pathogens in the *Candida/Clavispora* clade. Even with limited available strains outside of the human body, we find that these environmental strains are not closely related to each other, nor are they geographically clustered with their most closely related clinical isolates, pointing to the extreme versatility and wide distribution of these yeasts. From these data it is, therefore, hard to determine if the niche shift in these yeasts moved from specialized saprophyte to human pathogen as previously hypothesized [29]. A broad set of *Candida/Clavispora* samples detected with marker-based sequencing from diverse environments across the globe reinforces this notion [40]. Even though thermotolerance is a critical trait for fungi to survive in the human body [42], the expansion in distribution of these yeasts outside tropical regions challenges the hypothesis that global warming is a major evolutionary driver of pathogenicity in this clade [17, 29]. Additional evidence is needed to understand how selection on thermotolerant traits operates in yeasts from areas where the effects of global warming are less pronounced (average ambient temperature <37°C) and areas where these effects raise average ambient temperatures above 37°C. Therefore, it seems more plausible that these pathogens, found in harsh conditions such as seawater, already possessed traits that make them suitable human pathogens. The detection of these yeasts in humans might correspond with increases in at-risk populations over the past 30 years, for example changes in cancer, transplant, and HIV patient populations, ultimately representing a change in host environment [27]. This explanation is logically fits with what we know about the transmission dynamics of these yeasts in hospital settings, specifically their ability to exclusively cause severe disease or long-term asymptomatic carriage in heavily medicated (antibiotics, antifungal, immunosuppressants) patient populations with indwelling medical devices [31, 42].

As we continue to see an increase in vulnerable patient populations, evident by recent increases in intensive-care hospitalization due to COVID-19, we should be vigilant in our surveillance for yeasts belonging to this clade being isolated clinically from humans, and work to better understand how human-fungal interactions transmit fungal pathogens to these populations. Increased environmental sampling and obtaining isolates from these diverse sources will greatly enhance the phylogenomic dataset generated in this study and help to form a better picture of the evolutionary dynamics of the human pathogenic lifestyle in this clade.

## MATERIALS AND METHODS

### Taxon Sampling

A maximum of 20 isolates were randomly selected for species with more than one isolate available. A total of 108 isolates representing 18 different species were sampled, including 4 species newly sequenced in this study. These isolates were mostly obtained from clinical specimens (66.6%), followed by the environment (15.0%), plants (11.0%), animals (5.5%), and insects (0.9%) (S1 Appendix). Eight *Candida* spp. isolates (*C. haemulonii* CBS 5150, *C. heveicola* CBS 10701, *C. heveicola* CBS 7249, *C. dosseyi* CBS 10313, *C. duobushaemulonii* CBS 9754, *C. ruelliae* CBS 10815, *C. mogii* CBS 2032 and *C. hainanensis* CBS 10696) were obtained as lyophilized cultures from the Westerdijk Fungal Diversity Institute (https://wi.knaw.nl/) for whole genome sequencing (Table 1). We selected *Saccharomyces cerevisiae* and *Metschnikowia bicuspidata* as outgroups for our phylogeny. We rooted all trees on *S. cerevisiae*

### DNA Extraction and Whole Genome Sequencing

The eight lyophilized isolates were revived on Potato Dextrose Agar using aseptic technique. From each plate, a colony was randomly selected and used to inoculate 10 mL of Sabouraud Dextrose Broth in a 50 mL conical Falcon tube. The tubes were incubated at 25°C in a shaking incubator at 4000 rpm for 72 hours. After incubation, the cultures were centrifuged, and pellet was harvested. Cells were resuspended in a SDS buffer solution and vortexed. The contents were added to Qiagen PowerBead tubes with 0.5mm glass beads and vortexed for 5 minutes. The tubes were then heat shocked at 60°C for 60 seconds. DNA was extracted using Qiagen Blood and Tissue Extraction kit. The sample was eluted with warm sterile water rather than the supplied elution buffer. The ITS region was amplified (ITS1-F and ITS4) from the extracted DNA of each isolate and sequenced to confirm species identification and extraction quality. Library preparation was conducted at the University of Colorado Anschutz UCDAMC Genomics Core and sequenced on the Illumina NovaSeq 6000. Raw sequence read data are available in NCBI under BioProject PRJNA945431.

### Genome Assembly and Annotation

Workflows and scripts used to generate these data are publicly available on GitHub page: https://github.com/kyleschutz/candida_clavispora_workflow. Raw sequence reads from 82 isolates were de novo assembled in this study. Reads from 74 isolates were downloaded from the NCBI Sequence Read Archive using SRA Toolkit’s fasterq-dump. Reads from the 8 purchased isolates were also de novo assembled. Read quality was assessed with FastQC to inform read trimming parameters [44]. Raw reads were trimmed using Trimmomatic [45] with a sliding window cutoff of 4:10 and *de novo* assembled with SPAdes v. 3.15.2 [46]. Genome assemblies for 26 isolates without raw sequence data available were retrieved from NCBI’s GenBank. Assembly statistics were evaluated for all assemblies using Assemblathon 2 [47]. Given that we used a combination of *de novo* assemblies (n=82) and reference assemblies (n=26), we annotated all assemblies to quality control and standardize for downstream analyses using Funannotate version 1.8.9 [48]. Assembly data are available for download: https://doi.org/10.5281/zenodo.7742436.

To assess the quality of the assemblies and their subsequent annotations, *Candida haemulonii* B11899 was aligned against a high-quality reference genome of *Candida haemulonii* using a custom database query with BLASTN [49]. This validation test showed 100% identity between coding regions in the two assemblies. Assembly completeness was assessed using the BUSCO version 3 using Saccharomycetes OrthoDB v. 10 [50] for all strains.

### Identifying Orthologous Gene Clusters

From the 110 input genomes, nucleotide coding sequence data (CDS) were translated to amino acid sequences using EMBOSS v. 6.6.0 [51]. In parallel, CDS regions from all isolates were concatenated to serve as a reference for downstream translation back to nucleotide sequences. 619 single-copy orthologous genes were detected from translated CDS regions using Proteinortho6 and extracted using the grab_proteins.pl script [52]. Each gene cluster was individually aligned with MAFFT v. 6.240 [53]. In order to maximize the information available for phylogenetic analysis these closely related strains, the alignments were reverse-translated back to nucleotide data referencing the original CDS file with RevTrans Version 2.0 before trimming and gap removal [54]. The alignments were then trimmed using the “gappyout” parameter in trimAl [55].

### Phylogenetic Analysis

To prepare for the estimation of the maximum likelihood (ML) species tree, the trimmed CDS alignments were concatenated using catfasta2phyml.pl script (https://github.com/nylander/catfasta2phyml). The resulting alignment consisted of 1,152,540 alignment sites and 592,724 alignment patterns representing 110 input genomes. The ML tree was estimated with RAxML-NG v. 8.0 with the following parameters: GTR + GAMMA model, 100 bootstrap replicates.

Gene trees were estimated with RAxML-NG version 8.0 with each of the 619 CDS alignments using the General Time Reversible (GTR) + GAMMA model of nucleotide substitution, to allow for rate heterogeneity across sites. Support was assessed with 100 bootstrap replicates [56]. Individual ML gene trees are available at: https://doi.org/10.5281/zenodo.7742436.

A coalescent species tree estimation was conducted using the resulting gene trees in ASTRAL 5.7.1 [35]. Gene conflict was assessed by a conflict analysis of the ASTRAL coalescent species tree using PhyParts with a 50% bootstrap cutoff parameter [57]. Conflicting gene tree topologies were visualized with the phypartspiecharts.py script (https://github.com/mossmatters/MJPythonNotebooks/blob/master/PhyParts_PieCharts.ipynb). We also examined topological hypotheses by filtering individual gene trees for predicted relationships (e.g., clades of exclusively environmental samples). We carried out this filtering with PAUP* v. 4.0a169 [39].

## DATA AVAILABILITY

Workflows and scripts used to generate these data are publicly available on Kyle Schutz’s GitHub https://github.com/kyleschutz/candida_clavispora_workflow.

Raw read sequence data were deposited in NCBI’s SRA Database under BioProject PRJNA945431.

Genome assemblies and ML gene trees are available at: https://doi.org/10.5281/zenodo.7742436

## ACKNOWLEDGEMENTS

We thank Dr. Lara Vimercati for sharing DNA extraction protocols and Eli Paul for help DNA extraction. We also thank Dr. Ricardo Franco-Duarte for sharing DNA sequence data for *Clavispora santaluciae* strains.

This work utilized the RMACC Summit supercomputer, which is supported by the National Science Foundation (awards ACI-1532235 and ACI-1532236), the University of Colorado Boulder, and Colorado State University. The Summit supercomputer is a joint effort of the University of Colorado Boulder and Colorado State University.

## FIGURE CAPTIONS

**SUPPLEMENTAL Fig 1:**
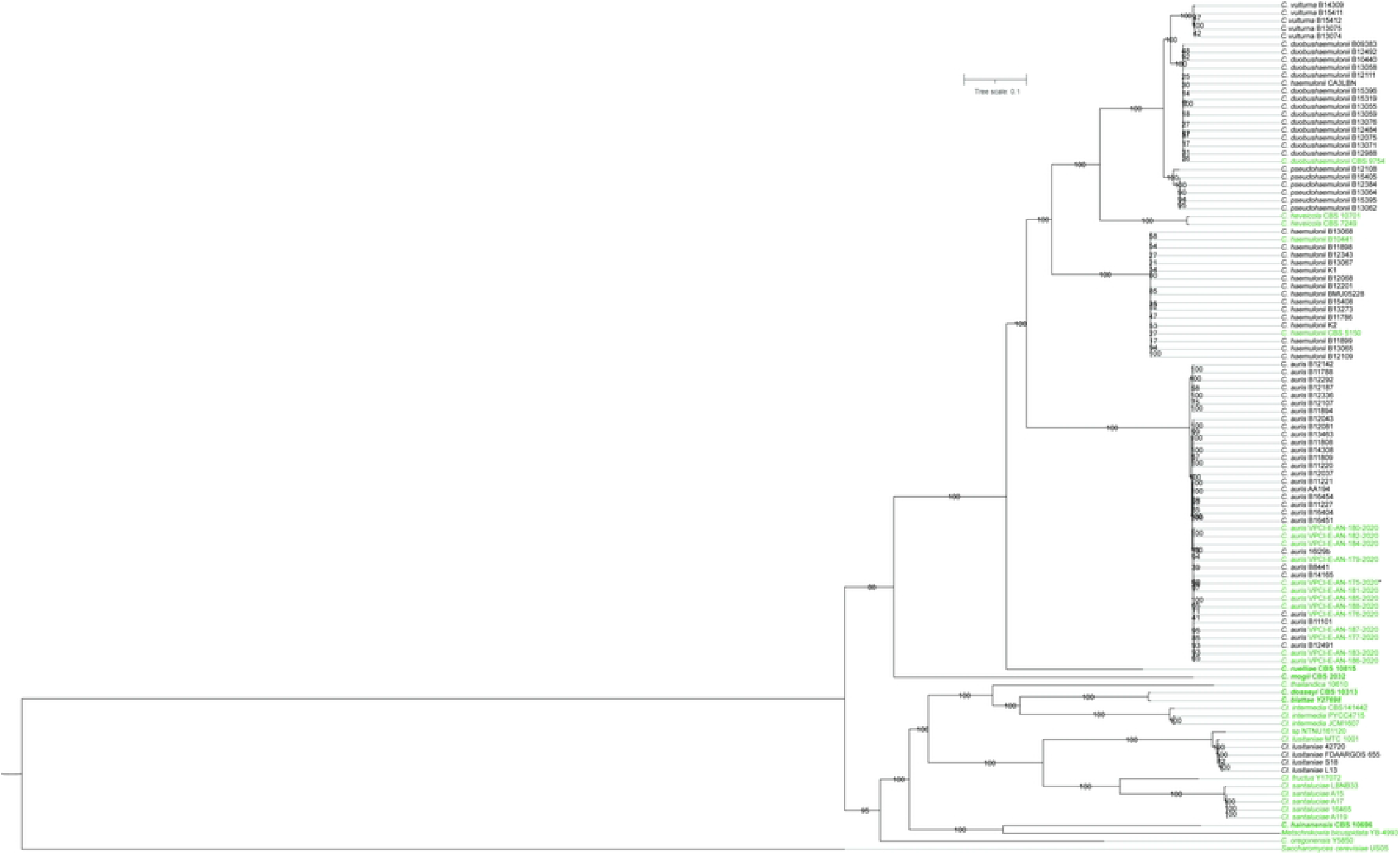
Expanded ML Phylogeny for *Candida/Clavispora*. A expanded maximum likelihood tree rooted on *S. cerevisiae* (pruned) based on 619 orthologous genes from 108 isolates using a GTR+GAMMA model and 100 bootstrap replicates. Bolded species had not been whole genome sequenced prior to this study. Green colored species indicate non-clinically isolated strains. The asterisk next to denotes an antifungal susceptible, environmentally isolated *C. auris* strain (fluconazole MIC, 8 mg/liter; amphotericin B MIC, 1 mg/liter) [29].

**SUPPLEMENTAL Fig 2:**
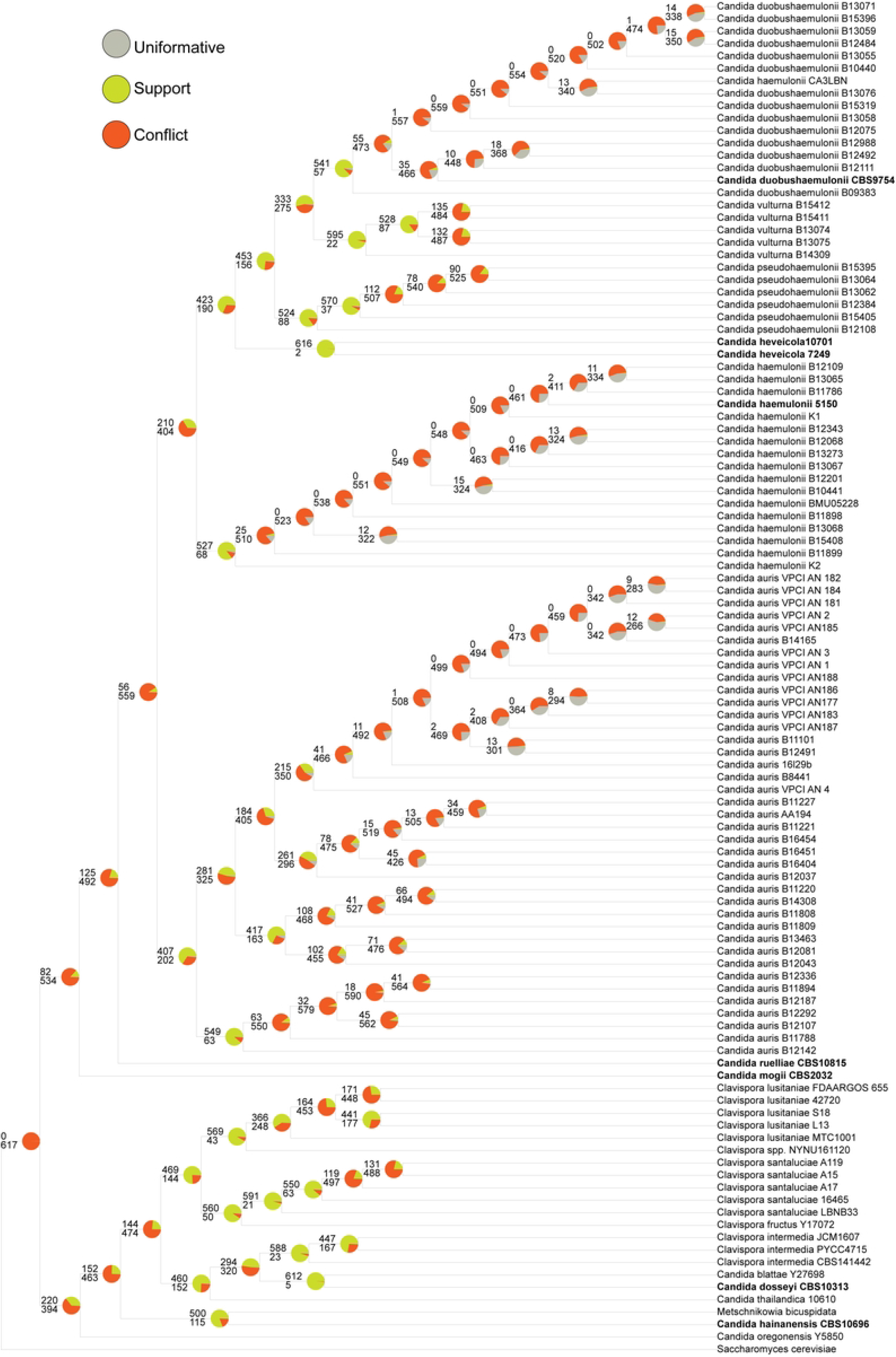
Gene conflict in *Candida/Clavsipora* yeasts. A coalescent species tree estimation based on 619 ML gene trees in ASTRAL. Gene conflict was assessed by a conflict analysis using PhyParts. Green portion of the pie charts show the proportion of 619 ML gene trees that support on the presented topology with 50% or higher bootstrap. Red portion of the pie charts show ML gene trees that support an alternate topology with 50% or higher bootstrap, while the grey portion of the pie charts depict genes that are uninformative (< 50% bootstrap support).

**S1 Appendix: Taxa metadata, including NCBI and SRA accession numbers**.

